# Metabolic Regulation of Single Synaptic Vesicle Exo- and Endocytosis in Hippocampal Synapses

**DOI:** 10.1101/2023.11.08.566236

**Authors:** Jongyun Myeong, Marion I Stunault, Vitaly A Klyachko, Ghazaleh Ashrafi

## Abstract

Glucose has long been considered a primary source of energy for synaptic function. However, it remains unclear under what conditions alternative fuels, such as lactate/pyruvate, contribute to powering synaptic transmission. By detecting individual release events in cultured hippocampal synapses, we found that mitochondrial ATP production from oxidation of lactate/pyruvate regulates basal vesicle release probability and release location within the active zone (AZ) evoked by single action potentials (APs). Mitochondrial inhibition shifted vesicle release closer to the AZ center, suggesting that the energetic barrier for vesicle release is lower in the AZ center that the periphery. Mitochondrial inhibition also altered the efficiency of single AP evoked vesicle retrieval by increasing occurrence of ultrafast endocytosis, while inhibition of glycolysis had no effect. Mitochondria are sparsely distributed along hippocampal axons and we found that nerve terminals containing mitochondria displayed enhanced vesicle release and reuptake during high-frequency trains, irrespective of whether neurons were supplied with glucose or lactate. Thus, synaptic terminals can entirely bypass glycolysis to robustly maintain the vesicle cycle using oxidative fuels in the absence of glucose. These observations further suggest that mitochondrial metabolic function not only regulates several fundamental features of synaptic transmission but may also contribute to modulation of short-term synaptic plasticity.

**Highlights:**

- Synapses can sustain neurotransmission across various activity levels by bypassing glycolysis and utilizing oxidative fuels.
- Mitochondria, but not glycolysis, regulate release probability and nanoscale organization of vesicle release within the active zone.
- Mitochondrial inhibition increases the occurrence of vesicle retrieval via ultra-fast endocytosis.
- Mitochondrial localization in nerve terminals enhances vesicle release and retrieval in the absence of glucose, representing a form of synaptic plasticity.

## INTRODUCTION

The brain requires a constant, ready source of energy to function properly. When the metabolic processes supporting energy generation go awry, for example secondary to hypoglycemia, ischemic stroke, or uncontrolled diabetes, loss of cognitive function follows rapidly^1,2^. Neurons, like many other cell types, convert carbon sources to ATP via glycolysis and oxidative phosphorylation (OXPHOS). Glucose has long been considered the primary carbon source for neurons in the mammalian brain^1^. However, glucose supply to the brain is both limited and subject to fluctuations^2–4^ thus placing neurons at risk of energetic stress. It has become increasingly clear that neurons can utilize several fuel types besides glucose, including lactate, which is supplied through the circulation or produced locally by astrocytes. Glucose undergoes glycolysis in the cytoplasm to produce ATP and pyruvate. Lactate, and its derivative pyruvate, are exclusively broken down via mitochondrial OXPHOS. The metabolic plasticity of neurons in utilizing multiple fuel sources is critical for their survival, yet how different energy sources impact neuronal communication remains poorly understood.

Nerve terminals are sites of metabolic vulnerability within neurons and rely on continuous energy supply to sustain neurotransmission. As such, terminals are heavily dependent on mitochondria for oxidative ATP synthesis and calcium buffering, as demonstrated by their heightened susceptibility to mitochondrial dysfunction^5^. Yet, mitochondrial distribution in neuronal axons is not uniform and it is estimated that 30-50% of hippocampal terminals lack mitochondria in their proximity^6–8^. Many studies have investigated the functional significance of mitochondria in nerve terminals but they have produced conflicting results. On one hand, it has been reported that the efficiency of neurotransmission does not vary significantly between terminals that do or do not contain mitochondria^9^. On the other hand, mitochondrial depletion has been shown to impair synaptic vesicle mobilization during sustained bouts of electrical activity, contributing to the variability in synaptic strength among nerve terminals^10–12^. Since hippocampal pyramidal neurons typically fire only short bursts of ∼3-25 action potentials (APs) at 1-20 Hz frequency^13,14^, these findings imply that mitochondrial function is largely dispensable for most physiological levels of activity in these neurons and is only required during intense firing. Thus, the question remains whether or not mitochondria regulate any aspects of synaptic transmission under physiological conditions.

The propagation of APs in synaptic boutons triggers exocytosis, reuptake/endocytosis, re-acidification, and re-docking of synaptic vesicles, collectively referred to as the synaptic vesicle cycle. Vesicle endocytosis is considered the most energetically susceptible step in the cycle, sustained by both glycolysis and mitochondrial OXPHOS following trains of AP firing^8^. Unlike endocytosis, the energetic demands of vesicle exocytosis are not well understood. Furthermore, metabolic regulation of the vesicle cycle has been primarily studied in response to trains of high-frequency stimulation rather than single APs. The resulting forms of neurotransmission are mechanistically distinct both in terms of release and reuptake. Specifically, in response to single AP firing, only docked vesicles belonging to the readily releasable pool (RRP) are released, while AP trains recruit vesicles from multiple pools, including the RRP, the recycling pool and, under some circumstances, even the reserve pool^15^. Moreover, under conditions of single AP firing, vesicles are primarily recovered through clathrin-independent fast or ultra-fast endocytosis^16^. In contrast, vesicles released during high-frequency AP trains are typically retrieved by the slower mode of clathrin-dependent endocytosis^16–18^. At present, it is not known to what extent glycolytic or mitochondrial ATP production powers single AP-driven vesicle release and retrieval.

Here, we investigated the energetics of neurotransmission in hippocampal terminals and uncovered that single AP-evoked vesicle release and endocytosis are regulated by mitochondria rather than glycolysis. Furthermore, we demonstrated that synaptic terminals can entirely bypass glycolysis to robustly maintain the vesicle cycle using oxidative fuels in the absence of glucose. We uncovered the physiological significance of mitochondrial localization in nerve terminals by showing that synapses without mitochondria exhibit slower and less efficient exocytosis as well as endocytosis compared to synapses that possess mitochondria. Our findings reveal the distinct energetic dependencies of single vesicle release and reuptake, underscoring the critical role of mitochondrial function in hippocampal terminals.

## RESULTS

### Mitochondria But Not Glycolysis Regulate the Spatial Properties Of Vesicle Release

Many steps in synaptic transmission are metabolically demanding, yet how these processes adapt to fuel availability is not well understood. We first examined the energetics of single AP-evoked vesicle release events in hippocampal synapses (**Figure 1A and B**) by selectively removing fuel sources or inhibiting their production pathways. Specifically, mitochondrial ATP production was acutely blocked using two complimentary agents (**Figure 1C**): by selectively inhibiting mitochondrial ATP synthase with oligomycin or by inhibiting mitochondrial pyruvate carrier (MPC) with UK5099. Glycolysis was inhibited by substituting glucose for lactate/pyruvate or using 2-deoxy-glucose (2DG), which serves as a competitive inhibitor of glucose metabolism via glycolysis. It has been previously shown that inhibition of either mitochondria or glycolysis leads to a reduction in ATP levels in nerve terminals^8^.

**Figure 1.**
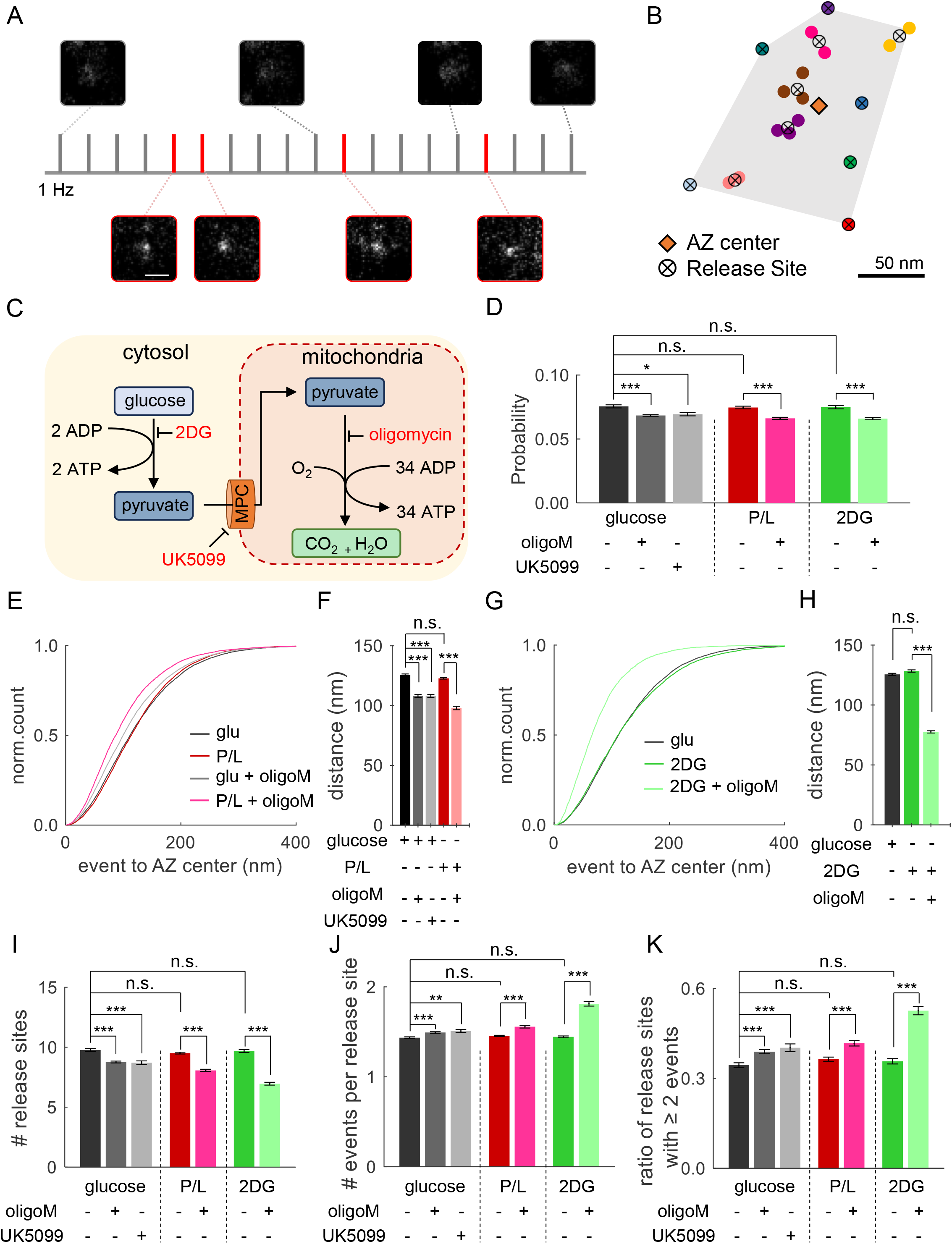
Mitochondrial ATP production regulates spatial properties of single release events. **(A)** Sample temporal distribution and representative images of single release events (red bars) or failures (gray bars) evoked by 1Hz stimulation in single boutons expressing vGlut1-pHluorin. Scale bar: 1 μm. **(B)** Sample hippocampal active zone (AZ) depicting localization of release events and release sites detected during recording. Events in the same release site are represented with the same color and cluster centers representing release sites are shown as crossed circles. **(C)** Schematic diagram of glucose metabolism for ATP production. Glycolytic breakdown of glucose within the cytoplasm forms pyruvate, yielding a total of 2 ATP molecules. Pyruvate enters mitochondria via the mitochondrial pyruvate carrier MPC and undergoes OXPHOS generating about 34 ATP molecules. Independently of glycolysis, pyruvate can also be derived from lactate, which is supplied through the circulation or produced locally by astrocytes. **(D)** Vesicle release probabilities were calculated for individual synapses stimulated at 1 Hz for 200 seconds under different metabolic conditions. OligoM and UK5099 are inhibitors of mitochondrial OXPHOS, while 2DG is a glycolytic inhibitor. **(E,G)** Cumulative distances of release events to the AZ center in synapses supplied with glucose, pyruvate/lactate, or 2DG with or without 2 μM oligoM. **(F,H)** Average distances of release events to the center of the AZ in data from (**E**) and (**G**). **(I)** Average number of clusters/release sites detected in individual AZs under different metabolic conditions. **(J)** Average number of events detected per release site under different metabolic conditions. **(K)** Ratio of release sites that had at least two events detected during the observation period to the total number of release sites. OligoM: oligomycin, 2DG: 2-deoxyglucose, P/L: equimolar mixture of pyruvate and lactate. ** < 0.01, ***p < 0.001, n.s., not significant. K-S test or One-way ANOVA.

To assess neurotransmitter release at a single-vesicle level, a pH-sensitive indicator pHluorin was targeted to the synaptic vesicle lumen via vGlut1 (vGlut1-pHluorin)^19,20^ using lentiviral infection at DIV3. To precisely localize individual vesicle release events within the synaptic active zone (AZ), we used a combination of near-TIRF imaging with well-established computational detection and localization approaches^21,22^. Individual release events were evoked by single AP stimulation at 1Hz at 37°C, recorded at 50 ms/frame for 120 s, and then automatically detected and localized with ∼27 nm precision, as previously described^23^ (**Figure 1A**). The AZ area was defined as the convolute hull of all events detected in an individual synapse, and its centroid was used to define the AZ center (**Figure 1B**). This “functional” definition of the AZ dimensions closely matches the ultrastructurally defined AZ dimensions observed by EM^23,24^. Hierarchical clustering of events with a 50 nm clustering diameter was then used to define individual release sites in each AZ^23–25^.

We first determined the basal vesicle release probability (Pr) at low activity levels (1Hz) and found that substituting glucose with pyruvate/lactate, or with 2DG, did not have any measurable effects on Pr (**Figure 1D**), suggesting that glycolysis is not required to maintain basal release. Inhibition of mitochondrial ATP production with either oligomycin or UK5099 resulted in significant reduction of Pr (**Figure 1D**), and the extent of reduction was the same whether glucose was present in solution or substituted with pyruvate/lactate (**Figure 1D**). This result suggests that synapses can efficiently utilize alternative fuels and that vesicle release evoked by single AP firing is more susceptible to the loss of mitochondrial ATP production than the loss of glycolysis.

Recent studies have shown that vesicle release is not randomly distributed across the AZ but has a precise spatio-temporal organization characterized by repeated utilization of several specialized release sites^21,22,24–28^. Because many steps in vesicle docking and release are ATP-dependent, we next asked whether the spatial organization of vesicle release exhibits metabolic plasticity and can adjust to different fuel types. We examined the spatial properties of the release events using the distance between each event and the AZ center as a simple criterion. As was the case with Pr, bypassing glycolysis by substituting glucose for pyruvate/lactate or using 2DG did not have any measurable effects on the spatial properties of vesicle release (**Figure 1E,F and 1G,H**). In contrast, mitochondrial inhibition (with oligomycin or UK5099) caused release events to occur significantly closer to the AZ center, and this effect of oligomycin was exacerbated by substituting glucose with pyruvate/lactate (**Figure 1E, F**) or with 2DG (**Figure 1G, H**). Moreover, in the presence of oligomycin, release events occurred at significantly shorter distance from each other than in control conditions (**Figure S1A**), and these intra-event distances were even shorter when oligomycin was combined with pyruvate/lactate or 2DG replacing glucose (**Figure S1A and B**). The shorter distances of the events to the AZ center as well as among the events themselves suggest a shift in release site utilization towards a subset of more centrally located sites. In support of this idea, we found that the number of release sites utilized by release events was significantly reduced in the presence of oligomycin or UK5099 (**Figure 1I**), while the number of events detected per release site was increased (**Figure 1J**). Accordingly, the number of repeatedly reused release sites was also increased (**Figure 1K**). As was the case in the earlier experiments, inhibition of glycolysis did not affect utilization of release sites (**Figure 1I-K**). We confirmed that the effect of oligomycin was not due to disruption of mitochondrial capacity for calcium buffering because GCaMP8f measurements of intra-terminal calcium revealed that none of our metabolic perturbations significantly altered the calcium rise amplitude or decay evoked by a single AP (**Figure S1E-I**). These results suggest that during single AP firing, spatial dynamics of vesicle release are sensitive to mitochondrial perturbations and that vesicle release shifts towards a subset of more centrally located release sites in the absence of mitochondrial OXPHOS. In contrast, vesicle release is largely insensitive to inhibition of glycolysis and can function normally by shifting to pyruvate/lactate-based mitochondrial ATP production.

### Mitochondria But Not Glycolysis Regulate Occurrence of Ultra-Fast Endocytosis

Vesicle exocytosis is rapidly followed by endocytosis, and various mechanistic and kinetically distinct forms of vesicle retrieval have been identified^29,30^. In our measurements, the kinetics of the pHluorin signal following vesicle fusion is believed to reflect, in large part, the endocytosis process^20,31^. Previous studies identified several kinetic components of endocytosis coupled to single-vesicle release events including an ultrafast endocytosis, with a decay time constant of ∼50-100 ms, and a fast endocytosis with a time constant on the order of 1s^29^. In agreement with the earlier findings, we observed two peaks in the Gaussian distribution of events’ decay kinetics (**Figure 2A, B**) with an average time course of ∼100 ms (23% of all events), and ∼500 ms (77% of all events), closely corresponding to ultra-fast and fast components of endocytosis, respectively. Whether different forms of single vesicle endocytosis exhibit distinct metabolic vulnerabilities remains poorly understood. We thus examined the decay kinetics of single AP-evoked vesicle release events in various metabolic conditions. Because of the noisy nature of single events, we used a mono-exponential fit to approximate decay kinetics for each event trace (**Figure 2A**); the resulting distribution of decay time constants was fit with two Gaussians to obtain the average fraction of ultra-fast and fast components of endocytosis in the synapse population and quantified as a ratio (**Figure 2B**). We found that glucose substitution with pyruvate/lactate or 2DG had no measurable effects on endocytosis kinetics of single events (**Figure 2C-E, S2A-D**). Specifically, the distribution of endocytosis kinetics across synapse population, as well as the ratio of the two components remained unaltered (**Figure 2C-E, S2A-D**). Thus, lack of glycolysis can be efficiently substituted by mitochondrial OXPHOS for the purposes of single vesicle endocytosis. While endocytosis also generally persisted after mitochondrial inhibition with oligomycin or UK5099, we noted an unexpected shift between the two forms of endocytosis across synaptic population with an increased prevalence in utilization of ultrafast endocytosis (**Figure 2D, 2E**). We obtained similar results when analyzing traces of evoked events averaged within individual synapses to mitigate the effects of noise (**Figure S2E-G**).

**Figure 2.**
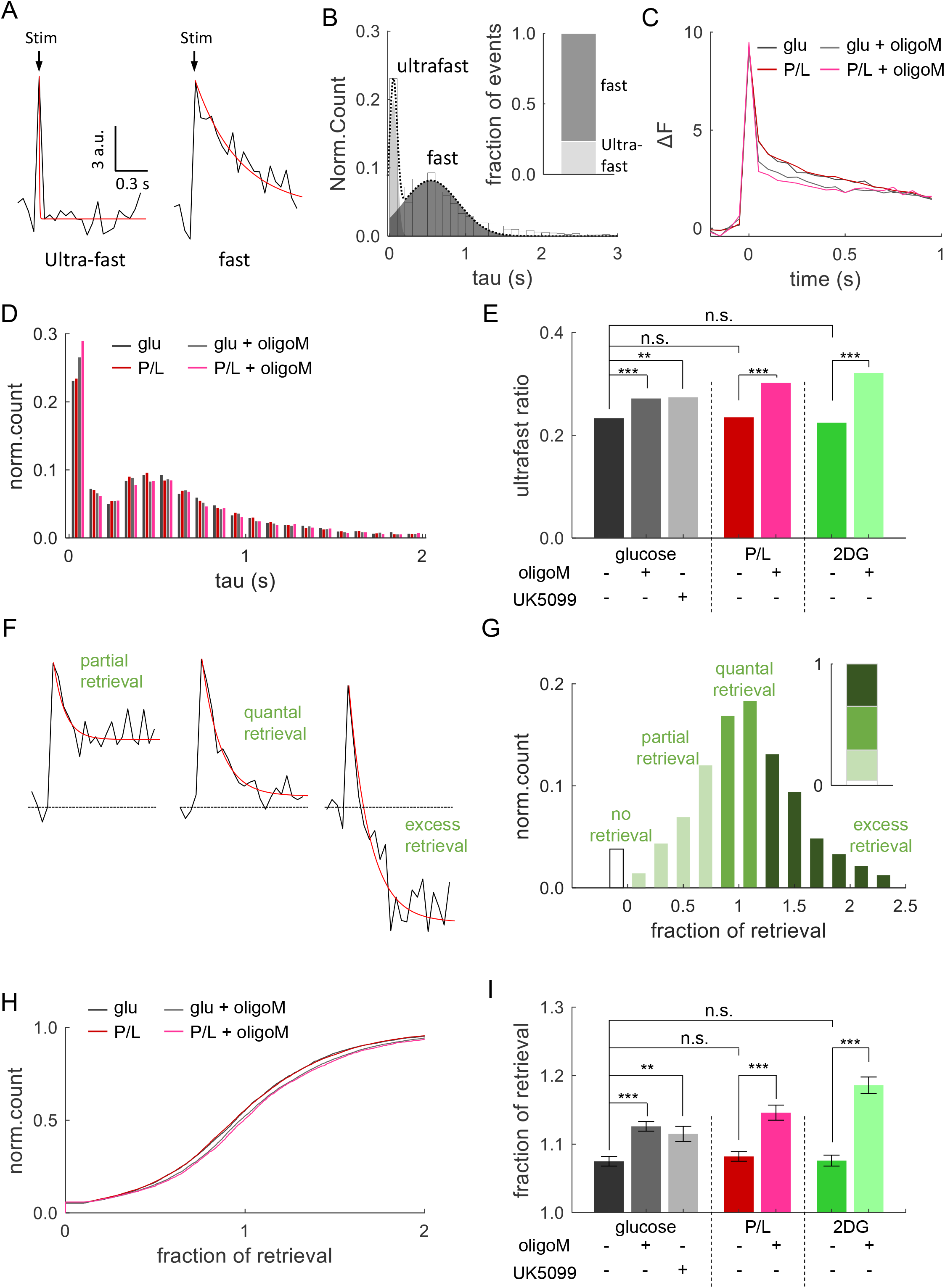
Mitochondrial inhibition increases the occurrence of ultrafast endocytosis. **(A)** Examples of single event traces showing two distinct rates of vesicle endocytosis following stimulation with 1 AP, characterized as ultrafast or fast. **(B)** The distribution of endocytosis tau values for 1AP-evoked release events. Dual Gaussian curves fitted to tau distribution for ultrafast (light gray) and fast endocytosis (dark gray). Inset: ratio of ultrafast to fast decay rates calculated based on the area under the tau distribution curve. **(C)** Average event traces recorded in glucose or pyruvate/lactate solutions with or without 2 μM oligoM. **(D)** Distributions of endocytosis tau values determined from fitting individual event decay with a mono-exponential fit in glucose or pyruvate/lactate solutions with or without 2 μM oligoM. **(E)** Average ratio of ultrafast endocytosis under different metabolic conditions. Statistical comparisons were based on comparing distributions of the corresponding tau values. **(F)** Representative examples of endocytic events displaying different fractions of retrieval with corresponding fitting traces (red lines). **(G)** Histogram of fractions of retrieval following single AP stimulation and its quantification into 3 groups (inset). **(H)** Cumulative plot of retrieval fractions in synapses supplied with glucose or pyruvate/lactate, with or without oligoM. **(I)** Average retrieval fractions under different metabolic conditions. *p<0.05, ** < 0.01, ***p < 0.001, n.s., not significant. K-S test or One-way ANOVA.

Endocytosis is characterized not only by decay kinetics, but also the timing of its initiation, commonly described using dwell time (**Figure S3A**). At the level of individual vesicles, the fluorescence dwell times detected after fusion represent the time during which pHluorin-tagged proteins remain at the presynaptic plasma membrane before retrieval. We observed that inhibition of neither glycolysis nor mitochondria significantly affected dwell times (**Figure S3B-D**), suggesting that initiation of endocytosis is governed by a relatively ATP-insensitive process. However, the half time, which is a broader measure of the overall duration of event decay, defined as the time from initiation to half maximum decay, was consistently shorter when mitochondria-mediated ATP production was inhibited with either oligomycin or UK5099 (**Figure S3E, F**). However, it was not affected by inhibition of glycolysis (**Figure S3E-G**), in agreement with our observations above.

The extent of endocytosis following single release events is known to vary widely within and across synapses, from partial/incomplete, to complete (i.e. quantal) and, in some cases, even excessive (**Figure 2F**)^32^. In our recordings ∼36% of the events exhibited a compete return to baseline, indicating full quantal retrieval of vGlut1-pHluorin, while ∼25% of fusion events showed partial retrieval, and ∼35% of events displayed excessive retrieval (**Figure 2G**). The remaining ∼4% did not show measurable retrieval during the 1 sec period between stimuli. We thus next examined if the extent of vesicle retrieval exhibits metabolic sensitivity. Fraction of retrieval was quantified as a ratio between the amplitude of the event decay and its peak amplitude, in which amount of event decay was determined as the amplitude difference between the peak and the final event amplitude (at 1 sec post stimulation). To take into account the noisy nature of the event traces, events with a ratio in the range of 0.8-1.2 were considered to be complete/quantal, events with ratio of less than 0.8 as incomplete, and >1.2 as excessive. Glucose substitution with pyruvate/lactate or 2DG had no measurable effects on the extent of retrieval, while mitochondrial inhibition resulted in a significant shift towards more excessive retrieval (**Figures 2H, 2I, and S2H, S2I**). Combined with the results above, these findings indicate that endocytosis coupled to single vesicle release is fully plastic in terms of fuel source, whether it be glucose or lactate/pyruvate. In contrast, mitochondrial inhibition alters endocytosis modes towards increased ultra-fast endocytosis and excessive retrieval, without changing the initiation of the endocytosis process.

Uni-vesicular (UVR) and multi-vesicular (MVR) are the two principal forms of evoked synchronous release. MVR refers to near simultaneous release of multiple (2 and more) vesicles and is ubiquitously observed in central synapses. In our measurements, MVR is commonly detected as the release of 2 vesicles and a corresponding second peak in the quantal event amplitude distribution (**Figure S4A and S4B**). We asked if the two forms of release differ in their metabolic sensitivity. If so, could a shift in UVR/MVR proportion explain the observed changes in vesicle release or retrieval in response to alterations in fuel availability? To examine this possibility, we performed detection of MVR events during 1Hz stimulation for 200 s and found that the probability of MVR was not significantly altered by any of our metabolic perturbations, including inhibition of glycolysis and mitochondrial OXPHOS (**Figure S4C**). Thus, changes in fuel availability did not significantly alter the occurrence of MVR and cannot explain the observed changes in synaptic transmission. We note that considering that the probability of MVR is ∼10 fold lower than that of UVR, it is still possible that small changes in MVR occurrence may exist but are too small to be detectable in our measurements.

We then examined endocytosis kinetics following MVR and noted that it had a much larger contribution from ultra-fast form compared to UVR in normal glucose conditions (UVR: 21%, MVR: 35%) (**Figure S4D, S4E**), which has been previously suggested based on simulations and ultrastructural studies^33^. As was the case with UVR, inhibition of glycolysis with 2DG had no effect on endocytosis kinetics following MVR events (**Figure S4I**), while addition of oligomycin shifted the ratio of the two components towards the ultra-fast component (**Figure S4F-I**), similarly to UVR (**Figure S4H-I**). We concluded that endocytosis coupled to two different forms of synchronous release have similar metabolic vulnerabilities, being fully functional in the absence of glycolysis, and exhibiting a similar shift toward ultra-fast endocytosis when mitochondrial ATP production was blocked.

### Mitochondria Sustain High-Frequency Synaptic Transmission in the Absence of Glucose

The above results suggest that both vesicle release and reuptake evoked by single APs exhibit high metabolic plasticity and can robustly utilize alternative fuels, such as lactate/pyruvate, while glycolysis is largely dispensable under these conditions. However, whether this metabolic plasticity can sustain synaptic transmission when energetic demands are increased during high-frequency stimulation is not well understood. We thus examined exocytosis and endocytosis evoked by high-frequency stimulation (40Hz, 50 AP) while altering fuel availability. Bypassing glycolysis by replacing glucose with pyruvate/lactate had no measurable effect on synaptic activity evoked by train stimulation, including both the rate of release and the rate of reuptake (**Figure 3A-C**). On the other hand, inhibition of mitochondrial ATP production with oligomycin or UK5099 significantly reduced both the rate of release and the rate of reuptake in the presence of glucose (**Figure 3A-F**), and nearly completely eliminated both processes when these agents were applied in media containing pyruvate/lactate replacing glucose (**Figure 3A-F)**.

**Figure 3.**
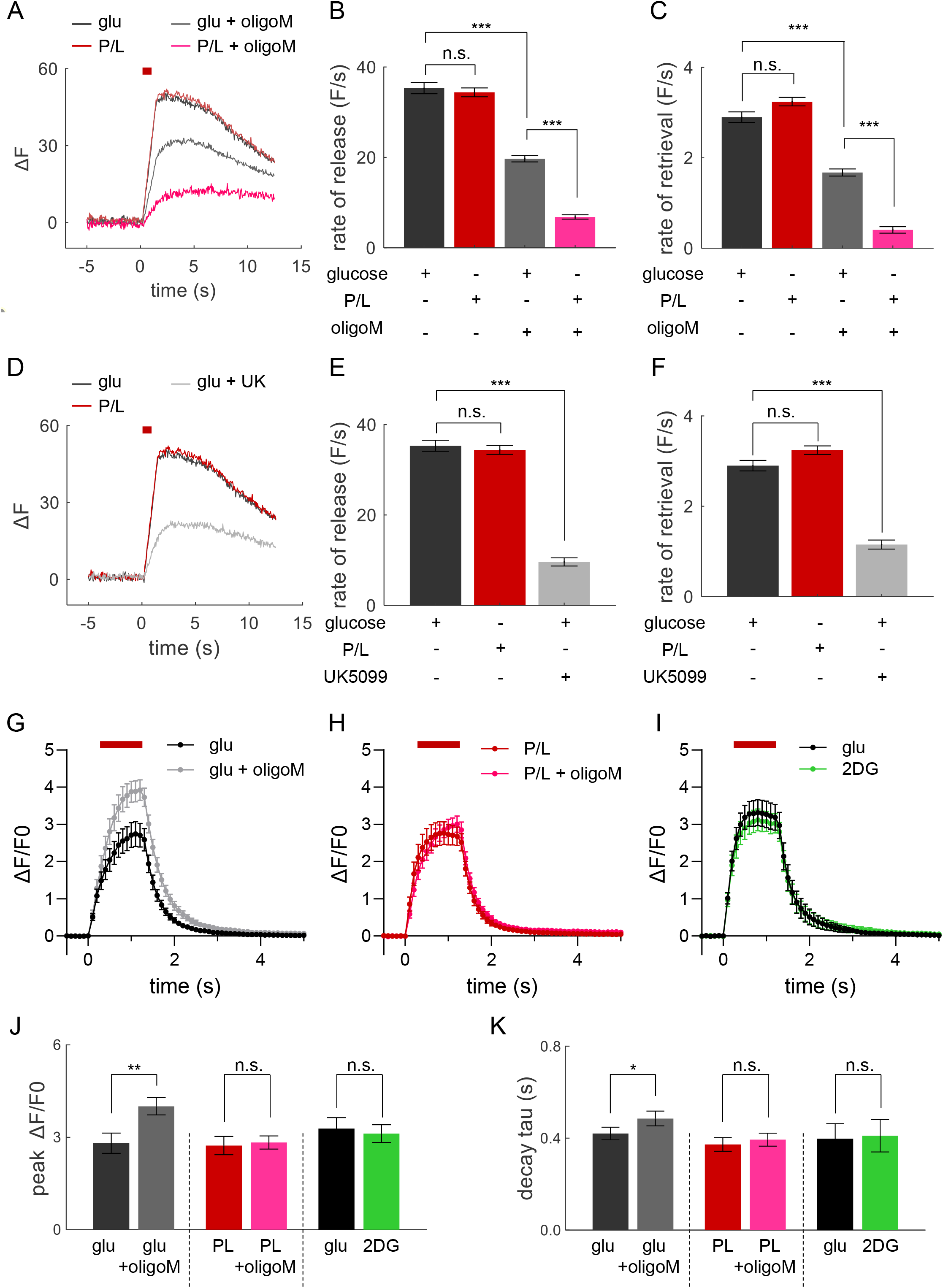
Mitochondria sustain high-frequency synaptic transmission in the absence of glucose. **(A)** Average vGlut1-pH fluorescence traces showing evoked release and reuptake during high-frequency stimulation. Neurons were incubated in solution containing glucose or pyruvate with or without 2 μM oligoM and stimulated with 50 AP at 40 Hz. **(B,C)** Vesicle exocytosis rates (**B**) and endocytosis rates (**C**) during the train stimulation in glucose or pyruvate/lactate solution with or without oligoM. **(D-F)** Same analysis as in (A-C), but with or without UK5099. **(G-I)** Intra-terminal calcium dynamics during train stimulation in neurons expressing jGCaMP8f in glucose, pyruvate/lactate, or 2DG, with or without oligoM. **(J,K)** Amplitudes (**J**) and decay constant (**K**) of calcium response during train stimulation under different metabolic conditions. *<0.05, ** < 0.01, ***p < 0.001, n.s., not significant. One-way ANOVA or paired t-test.

Since both exo- and endocytosis are calcium-dependent, and neurotransmitter release undergoes calcium-driven facilitation during high-frequency trains, we further examined if the observed changes in synaptic transmission are caused by disruption of presynaptic calcium dynamics that may arise if mitochondrial calcium buffering capacity is altered by our metabolic perturbations. Using GCaMP8f measurements, we observed that calcium rise and decay remained unchanged in all metabolic conditions used (**Figure 3G-K**) except for oligomycin, which caused a slightly larger calcium elevation during trains (**Figure 3G,J, and K**). Notably, this effect of oligomycin on calcium elevation is opposite in direction of change to the reduction of vesicle release caused by oligomycin, and thus cannot serve as the mechanistic basis for this observation. Thus, we conclude that the observed effects of mitochondria inhibition on synaptic transmission is mediated primarily by alterations in mitochondrial ATP production rather than its calcium buffering capacity.

Together, these results indicate that hippocampal synapses exhibit strong metabolic plasticity and can efficiently adapt to changes in fuel sources even during high-frequency activity, but are more susceptible to loss of mitochondrial ATP production than loss of glycolysis. The distinctive effects of mitochondrial inhibition on endocytosis during trains versus single release events also suggest that different forms of endocytosis are utilized following these stimulation paradigms, with a shift from fast and ultra-fast forms during single AP stimulation towards bulk endocytosis during elevated activity^34^.

### Mitochondrial Presence at the Synapse Enhances Exo- and Endocytosis During High-Frequency Stimulation

What are the functional implications of these findings to synaptic activity across different nerve terminals? Previous studies indicate that fuel availability at the synapses is highly variable even under normal conditions. In particular, only a proportion (∼50-70%) of synapses contain mitochondria^6–8^, and in adult neurons most of mitochondria are largely stationary on millisecond-to-seconds timescales^35^. This limits mitochondria’s ability to supply ATP to distant synaptic boutons during bouts of high-frequency activity, which typically last only a few seconds in hippocampal excitatory neurons^13^. Furthermore, it is unclear if ATP supply from glycolysis is sufficient to fully compensate for mitochondrial absence. Therefore, we investigated how presence or absence of mitochondria at the synapse affects neurotransmitter release during high-frequency activity. Mitochondria were localized using red-shifted MitoTracker (**Figure 4A, B**) and a threshold was placed on mitochondrial fluorescence intensity to reliably distinguish it from the axonal background (**Figure 4C**). Synaptic boutons were considered to be directly associated with mitochondria if the distance from the bouton center to the nearest mitochondrion edge was <1µm (**Figure 4B**). By this definition, 69% (358/518) of synapses were directly associated with a mitochondrion in our culture conditions, while 31% (160/518) of synapses were without mitochondria. Tracking mitochondria movements revealed that only ∼2% of mitochondria exhibited fast mobility (speed>0.5µm/s), while the vast majority (∼80%) were nearly immobile or exhibited very slow mobility (speed<0.1µm/s) (**Figure 4D**). We then compared synaptic responses to high-frequency stimulus trains (50APs at 40Hz) between synapses with and without mitochondria. While both subpopulations of synapses had a robust response to high-frequency stimulus trains (**Figure 4E**), the response amplitude and the corresponding rate of vesicle release were significantly larger in synapses with mitochondria than in those without (**Figure 4E, 4F**). Moreover, the rate of endocytosis following release was significantly faster in synapses with mitochondria than in those without mitochondria (**Figure 4G**). To confirm that these differences were dependent on mitochondria, we acutely blocked mitochondria ATP production with oligomycin which reduced synaptic response selectively in synapses containing mitochondria (**Figure 4E**). Oligomycin also eliminated differences in the rate of release and the rate of endocytosis between synapses with and without mitochondria (**Figure 4F, G**), thus confirming that the observed effects are due to the presence of mitochondria at synapses. When oligomycin was applied in the presence of lactate/pyruvate replacing glucose, it eliminated release almost entirely (**Figure 4E-G**), supporting the notion that there are no other major sources of ATP remaining at the synapses under these conditions. In summary, our observations suggest that availability of mitochondria at the synapse regulates the efficiency of synaptic vesicle release and uptake, thus representing a form of mitochondrially-driven synaptic plasticity.

**Figure 4.**
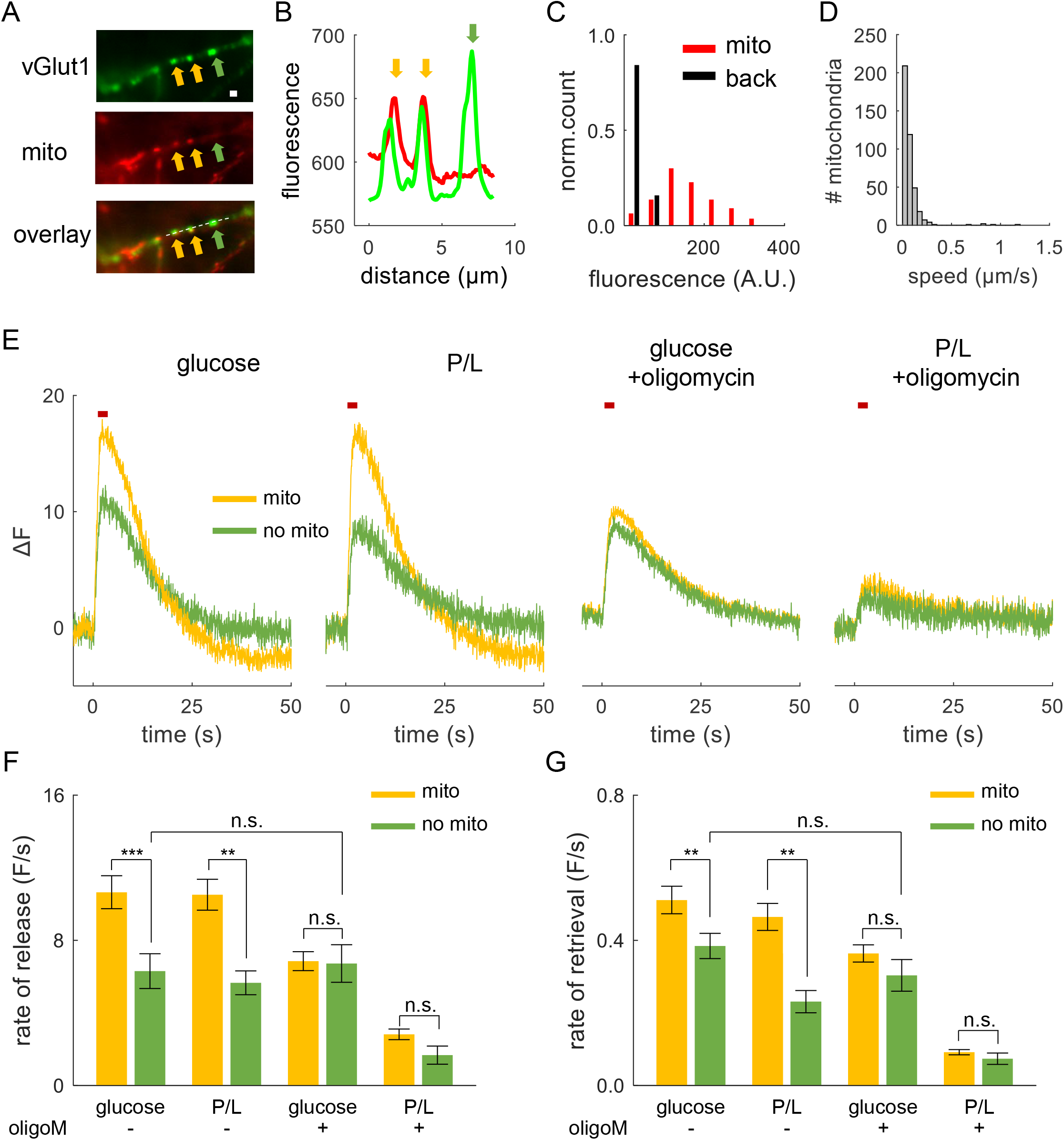
Mitochondrial presence at the synapse enhances exo- and endocytosis during high-frequency stimulation. **(A)** Sample combined fluorescence images (averaging all frames in each recording) of an axon showing synaptic boutons (vGlut1-pH, upper, green), the mitochondria (MitoTracker, middle, red), and overlay of the two images (bottom). The arrowheads indicate examples of synapses associated with mitochondria (yellow) or those without mitochondria (green). Scale bar = 1 μm. **(B)** Line scans of vGlut1-pH and MitoTracker along the dashed line in (**A**) with the corresponding arrows. **(C)** Distribution of background (black) and mitochondrial fluorescence intensities (red) in hippocampal axons labeled with MitoTracker. The distribution was used to define the threshold for robust mitochondria detection above background. **(D)** Distribution of mitochondrial motility speeds determined over 10s of imaging. **(E)** Traces of vGlut1-pH responses to 40Hz 50 AP simulation in synapses with (yellow) or without (green) mitochondria in glucose or pyruvate/lactate solution, with or without oligoM. The red bar represents the stimulation period. **(F)** Vesicle release rates in synapses with or without mitochondria during train stimulation under different metabolic conditions. **(G)** Vesicle retrieval rates measured as the rate of fluorescence decline 5-15 s after the end of stimulation under different metabolic conditions. **p<0.01, ***p < 0.001, n.s., not significant. Two-sample t-test.

## DISCUSSION

Glucose has long been considered the primary energy source for neurons, but accumulating evidence suggests that neurons can also use other fuels such as lactate and pyruvate, which are present in the circulation and can cross the blood-brain barrier^36^, or are locally supplied to neurons by neighboring astrocytes^37^. Lactate and pyruvate are strictly oxidative fuels that unlike glucose, cannot undergo glycolysis. To what extent these alternative fuel types are utilized during synaptic transmission is not well understood. Here, we demonstrate that lactate/pyruvate can fully replace glucose in supporting vesicle release and retrieval evoked by single APs as well as by high-frequency trains in hippocampal excitatory synapses. Therefore, our findings indicate that synaptic transmission exhibits extensive metabolic plasticity and can function robustly in the absence of glucose, with lactate/pyruvate as the main metabolic fuel. The remarkable metabolic plasticity of synaptic transmission in utilizing multiple fuel sources is critical for sustaining cognitive function amidst fluctuating glucose availability in the brain that normally occurs in various physiological conditions such as sleep or fasting. Importantly, our results demonstrate that mitochondrial energy production contributes to setting several fundamental properties of synaptic transmission, including the release probability, localization of vesicle fusion, and the kinetics of vesicle retrieval. We determined that mitochondrial function has important implications for rapid forms of synaptic plasticity since localization of mitochondria at the synapses enhances both vesicle exocytosis and retrieval, regardless of whether neurons are supplied with glucose or lactate. Thus, mitochondrial ATP production not only supports fundamental aspects of synaptic transmission during single AP firing and high-frequency activity, it also represents a mechanism contributing to the variability of synaptic responses and short-term synaptic plasticity.

Hippocampal synapses transmit information across a range of frequencies and durations. Different modes of transmission are expected to have distinct energetic requirements, particularly with respect to their dependence on mitochondrial and glycolytic ATP production. Due to its capacity for rapid ATP production, it was thought that glycolysis powers synaptic transmission at basal activity levels, with mitochondrial OXPHOS coming into play only during high-frequency trains^38,39^. In contrast, our results suggest entirely different roles for these two main energy pathways in supporting synaptic transmission. Specifically, we observed that mitochondrial ATP production is not only necessary to sustain robust vesicle release in response to high-frequency trains, but it is also indispensable for vesicle release evoked by single APs. In contrast, both vesicle release and reuptake following single APs are unaffected by lack of glycolysis. These findings raise the question: under what conditions glycolysis becomes indispensable to the energetics of synaptic transmission? Our findings collectively suggest that glycolysis only becomes a critical source of ATP for the vesicle cycle during prolonged firing which we previously showed leads to the activation of the energy sensor AMP kinase and recruitment of the glucose transporter GLUT4 to the presynaptic surface^40^. Notably, our findings do not imply that mitochondriaexclusively power synaptic transmission without contribution from glycolysis,. Rather, our results can be interpreted to indicate that ATP derived from both pathways is normally used to power basic synaptic processes, but there is a stronger dependence on mitochondrial ATP production. While this may appear counterintuitive at the first glance, the ATP yield of OXPHOS (32 molecules per glucose) is considerably higher than glycolysis (2 molecules per glucose) and it can be rapidly upregulated by calcium uptake into the mitochondrial matrix^41^, thus fine-tuning ATP production to activity levels. Despite its lower yield, glycolysis can be rapidly localized to sites of high ATP consumption through compartmentalization of its enzymes^42,43^ to provide additional ATP for high-frequency trains.

Our data demonstrate that mitochondrial ATP, but not glycolysis, is necessary to sustain several key features of neurotransmitter release evoked by single APs, including both the basal release probability and the proper localization of vesicle release within the AZ. Indeed, inhibition of mitochondrial ATP production not only reduced the Pr overall, but it also shifted vesicle release closer to the center of the AZ suggesting that the energetic barrier for vesicle release is lower in the AZ center that the periphery. This observation provides a mechanistic basis for our earlier findings that more centrally located release sites have higher utilization rates and thus higher release probability at low activity levels^23^. This energetic barrier can be overcome during high-frequency stimulation, presumably due to the elevation of intra-terminal calcium resulting in the engagement of more peripheral release sites^23^. It remains to be determined why the energetic cost for vesicle release near the AZ center is lower than at the periphery. However, it is notable that the nanoclusters containing components of the presynaptic machinery for vesicle docking and release are larger and more stable at AZ center^44^. Therefore, release sites near the AZ center may not require as much *de novo* ATP synthesis, in contrast to the AZ periphery where the presynaptic machinery is more mobile and may require more energy for capture/stabilization^28,45–48^. This spatial pattern of metabolic sensitivity may be also attributed to localized ATP gradients within the AZ which would be difficult to examine with the existing optical ATP sensors. Interestingly, we found that inhibition of mitochondrial ATP production produced stronger effects on the location and probability of vesicle release in the presence of 2DG than in lactate/pyruvate. This is likely because 2DG not only inhibits glycolytic ATP production, but also reduces the availability of pyruvate for OXPHOS, which further highlights the energetic dependency of single vesicle release. Interestingly, some vesicle release at the AZ center persisted when both glycolysis and OXPHOS were blocked with 2DG and oligomycin, suggesting that the steady-state concentration of ATP, estimated to be ∼1.5mM^49^, may be sufficient to power release during low levels of activity.

Vesicle retrieval is known to be an energetically demanding process^9,49^. Our results suggest that fuel availability, and specifically mitochondrial ATP, is an important factor that determines, at least in part, the utilization of different forms of vesicle retrieval. Following single AP stimulation, vesicle reuptake is primarily mediated by clathrin-independent fast endocytosis and ultrafast endocytosis^29^, which in our measurements are coupled to ∼77% and ∼23% of release events, respectively. Surprisingly, we found that mitochondrial inhibition resulted in increased occurrence of ultrafast endocytosis, both following UVR and MVR, suggesting that this form of endocytosis might have a lower energetic barrier. This is especially intriguing in light of a recent study reporting that Dynamin 1 molecules are pre-recruited to endocytic sites for ultrafast endocytosis^50^ and this very rapid process is driven by membrane compression which propagates along the membrane surface with very low energy demand^33^. In line with the increased engagement of ultra-fast endocytosis, we observed overall faster pHluorin signal recovery/shorter event half-time and an increased prevalence of complete/excessive endocytosis when mitochondrial ATP production was blocked. Interestingly, the timing of initiation of vesicle retrieval appears to be largely unaffected by inhibition of either or both of the main ATP production pathways. This suggests that the energetic demand for initiation of endocytosis is sufficiently low for it to be powered by residual ATP in nerve terminals when both pathways are inhibited. Notably, in contrast with single AP stimulation, endocytosis following high-frequency trains was significantly slower when mitochondrial ATP production was inhibited. These differences are in line with previous studies showing that the preferential mode of endocytosis shifts from fast or ultra-fast endocytosis following single APs to bulk endocytosis during high-frequency trains^34^, which presumably has higher energetic costs.

The central role of mitochondria in the metabolic support of vesicle release and retrieval is particularly intriguing considering that, under physiological conditions, mitochondria are sparsely distributed in hippocampal axons, leading to a significant portion of terminals lacking these organelles at any given point in time. Indeed, we found that mitochondrial localization in nerve terminals enhances efficiency of both vesicle release and reuptake, irrespective of whether neurons are supplied with glucose or lactate/pyruvate. We attribute these mitochondrial effects to its role in ATP production rather than in calcium buffering since changes in intra-terminal calcium during mitochondrial inhibition could not account for reduced release probability. Mitochondria are also involved in the production and buffering of reactive oxygen species (ROS), however ROS signaling has only been implicated in long-term synaptic plasticity^51–56^, and not short-term plasticity. Thus, our study implicates mitochondria as a critical player in the short-term plasticity of synaptic transmission and the heterogeneity of synaptic response across nerve terminals. These findings add a new dimension to the plethora of the mechanisms that modulate synaptic plasticity on the timescales of seconds, which is relevant to many critical information processing functions including working memory and decision making^57,58^.

The mitochondrial network in axons is not static; rather it is dynamically modulated by many signaling mechanisms including release of serotonin, dopamine, and glutamate spillover, which control mitochondrial transport, fission and fusion, as well as local attachment to actin or microtubules^59–61^. Considering that the typical speed of axonal mitochondrial movement is on the order of ∼10um per minute^59^, mitochondrial trafficking and the remodeling of the mitochondrial network in relation to synapses could function as a mechanism to modulate synaptic function in response to energetic stress. Indeed, activation of the energy sensor AMP kinase has been shown to drive the recruitment and anchoring of mitochondria in presynaptic terminals. In future, it will be important to investigate how prolonged exposure to various neuronal fuels like glucose, lactate, or ketone bodies impacts the localization of mitochondria within nerve terminals, and what functional implications this holds for synaptic transmission. Understanding mitochondrial function in the synapse will also shed light on how defects in mitochondrial motility, fission and fusion, and metabolic function contribute to the pathogenesis of neurodegenerative diseases, such as, amyotrophic lateral sclerosis, Parkinson’s, and Alzheimer disease (AD).

## ACKNOWLEDGEMENTS

We thank M. Laramie for preparation of primary hippocampal cultures, and the Hope Center Viral Vectors Core at Washington University, Saint Louis, MO. This work was funded by NINDS R35 NS111596 (V.K.), NIGMS R35GM147222 (G.A.), and the Whitehall Foundation (G.A.). The authors declare no competing financial interests.

## Author contributions

Jongyun Myeong: Investigation, visualization, formal analysis, and writing (original draft, review, and editing). Marion I. Stunault: Investigation, visualization, formal analysis, and writing (original draft, review, and editing). Ghazaleh Ashrafi: Conceptualization, funding acquisition, supervision, and writing (original draft, review, and editing). Vitaly Klyachko: Conceptualization, funding acquisition, supervision, and writing (original draft, review, and editing).

## STAR★METHODS

### KEY RESOURCE TABLE

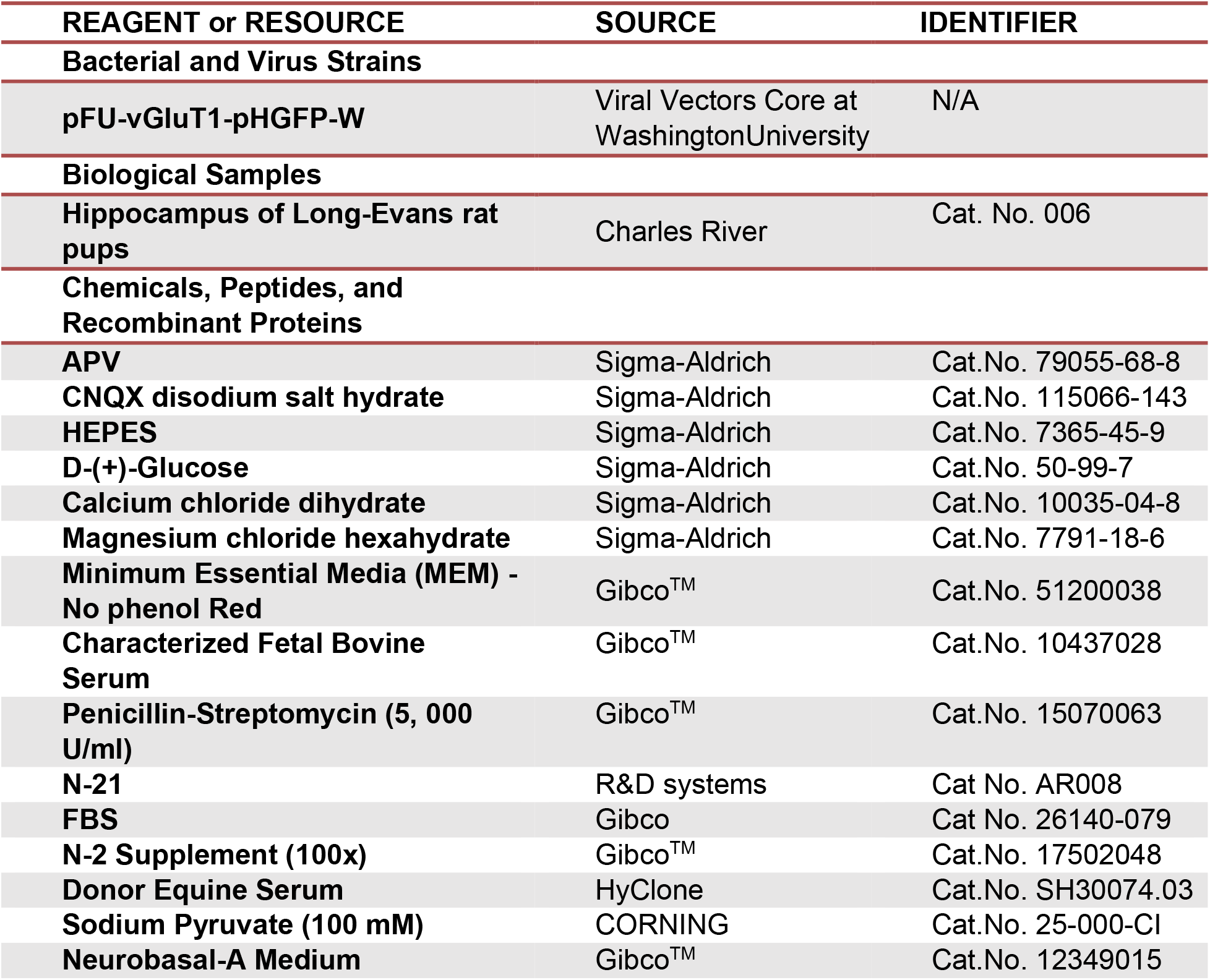
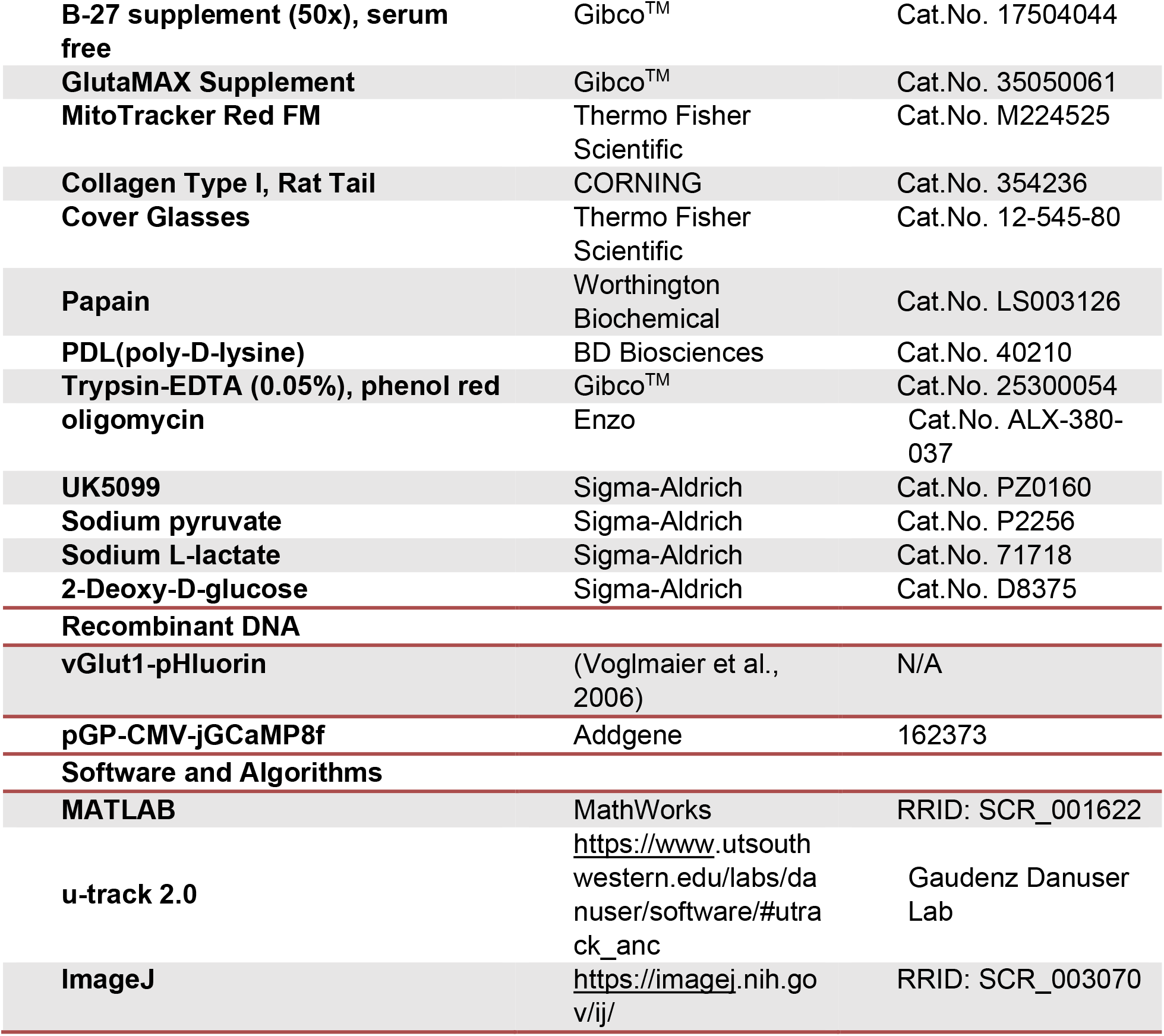

### RESOURCE AVAILABILITY

#### Lead contact

Further information and requests for resources and reagents should be directed to and will be fulfilled by the Lead Contacts, Dr. Vitaly A. Klyachko and Dr. Ghazaleh Ashrafi

#### Materials availability

This study did not generate new or unique reagents or other materials.

#### Data and code availability

- This paper does not report standardized data types. All data reported in this paper will be shared by the lead contacts upon request.
- This paper does not report standalone custom code. MATLAB was used to appropriately organize, process, and analyze data and corresponding routines are available from the lead contacts upon request.
- Any additional information required to reanalyze the data reported in this paper is available from the lead contacts upon request.

### EXPERIMENTAL MODEL AND SUBJECT DETAILS

#### Animals

All animal procedures conformed to the guidelines approved by the Washington University Animal Studies Committee. Wild-type rats of the Sprague-Dawley strain were used for preparation of primary neuronal cultures.

#### Neuronal Cell Culture

Neuronal cultures were produced from the hippocampus of rat pups of mixed gender as previously described^22,23,41^. Briefly, the hippocampal were dissected from E16-17 pups, dissociated by papain digestion, and plated on glass coverslips. Neurons were cultured in Neurobasal media supplemented with B-27 supplement. Neurons were grown on top of a confluent astrocyte monolayer. For calcium imaging experiments, hippocampi were dissected from 1-3 days old rat pups of mixed sex. Tissues were dissociated and mixed cultures of neurons and glia were plated on coverslips coated with poly-ornithine. Hippocampal neurons were maintained in culture media composed of MEM (Thermo Fisher Scientific 51200038), 0.6% glucose, 0.1 g/L bovine transferrin (Millipore 616420), 0.25 g/L insulin, 0.3 g/L GlutaMAX^TM^ supplement (Thermofisher 35050-061), 5% fetal bovine serum (R&D Systems S11510), 2% N-21 (R&D Systems AR008).

In all cultures, 4 μm cytosine β-d-arabinofuranoside (Ara-C) was added after 2-3 days in vitro (DIV) to limit glial proliferation. Cultures were incubated at 37°C in a 95% air and 5% CO2 humidified incubator for 14 - 16 DIV prior to use.

### METHOD DETAILS

#### Neuronal Transfection*/*Lentiviral Infection

VGluT1-pHluorin was generously provided by Drs. Robert Edwards and Susan Voglmaier (UCSF). Lentiviral vectors were generated by the Viral Vectors Core at Washington University. Hippocampal neuronal cultures were infected at DIV3, as previously described^22,23^. For calcium imaging, neurons 5-7 DIV were transfected with the pGP-CMV-jGCaMP8f plasmid using the calcium phosphate method.

#### Microscopy

*Near-TIRF microscopy*. Experiments were conducted at 37°C within a whole-microscope incubator chamber (TOKAI HIT). vGlut1-pHluorin was excited with a 488 laser (Cell CMR-LAS-488, Olympus) or with 475 nm LED (Olympus) for near-TRIF or wide-field imaging and MITO tracker Red FM was excited by 575 nm LED. Fluorescence was monitored using an inverted microscope (IX83, Olympus) and a 150x/1.45NA objective. The Z-drift compensation system (IX3-ZDC) was used to ensure a constant position of the focal plane during imaging. Near-TIRF was achieved by adjusting the incident angle to 63.7°, which is near the critical angle of 63.63°. Images were acquired every 50 ms (with an exposure time of 49.38ms) using a cooled EMCCD camera (iXon life 888, ANDOR). Field simulation was performed by using a pair of platinum electrodes and controlled by the software via Master-9 stimulus generator (A.M.P.I.). Samples were perfused with bath solution (125 mM NaCl, 2.5 mM KCl, 2mM CaCl_2_, 1mM MgCl_2_, 10 mM HEPES, 30 mM Glucose, 50 μM APV, 10 μM CNQX, adjusted to pH 7.4). In some experiments glucose was substituted for 1.5 mM lactate and 1.5 mM pyruvate, or 30 mM 2DG, as specified in the text.

#### Calcium imaging

Experiments were performed on a custom-built laser illuminated epifluorescence microscope with an Andor iXon Ultra 897 camera. Coverslips were mounted in a laminar flow perfusion chamber and perfused with Tyrodes buffer containing (in mM) 119 NaCl, 2.5 KCl, 2 CaCl2, 2 MgCl2, 50 <u>HEPES</u> (pH 7.4), 5 glucose or 1.5 lactate and 1.5 pyruvate, supplemented with 10 μM 6-cyano-7nitroquinoxalibe-2, 3-dione (CNQX), and 50 μM D,L-2-amino-5phosphonovaleric acid (APV) to inhibit post-synaptic responses. APs were evoked in neurons with 1 ms pulses creating field potentials of ∼10 V/cm via platinum–iridium electrodes. Temperature was maintained at 37°C using an Okolab stage top incubator. Cytosolic calcium signaling in axons of hippocampal neurons was measured using the GCamp8f sensor as previously described^62^. Neurons were stimulated with trains of 50 APs at 40Hz or 10APs at 1Hz. As a standard, 50 frames were recorded before the stimulus train was triggered.

#### Pharmacology

UK5099 and oligomycin were diluted in dimethyl sulfoxide (DMSO) and stored at 4°C and - 20°C respectively. Samples were pre-incubated in imaging solution with 2 µM of oligomycin, or 50 µM of UK5099, for 5-10 minutes prior to the beginning of the recordings. The effective final DMSO concentration was <0.5%. 30 mM 2DG or 1.5mM lactate and 1.5mM pyruvate were used as alternative fuels, respectively.

### QUANTIFICATION AND STATISTICAL ANALYSES

#### Event detection and localization using mixture-model fitting

VGluT1-pHluorin–based release event detection and localization at subpixel resolution were performed as previously described^21–23^ using Matlab and the uTrack software package, which was kindly made available by Dr.Gaudenz Danuser’s lab. Localization precision was determined from least-squares Gaussian fits of individual events as previously described^23^. To determine the basal Pr, boutons were stimulated at 1 Hz and Pr was determined by calculating the probability of detecting release events as determined over a 200-second period.

#### Detection of MVR

MVR was determined by calculating the mean and standard deviation values of peak intensities from individual synapses during 1Hz 200 sec stimulation period. Events with transient intensities exceeding the sum of the mean and twice the standard deviation were designated as MVR, while the remaining events were classified as UVR.

#### Definition of AZ dimensions and center

The AZ size was approximated based on the convex hull encompassing all vesicle fusion events in a given bouton and its centroid was used to define the AZ center. This measurement is in close agreement with the ultrastructural measurements of AZ dimensions^23^.

#### Definition of release sites

Release sites were defined using hierarchical clustering algorithm with a clustering diameter of 50nm using built-in functions in Matlab as described^23–25^. We have previously shown that the observed clusters do not arise from random distribution of release events, but rather represent a set of defined and repeatedly reused release sites within the AZs.

#### Analysis of endocytosis

The ratio of ultra-fast/fast endocytosis, which we refer to as ultrafast ratio, was defined by comparing the areas of the Gaussian fits to endocytosis tau histogram of fast vs ultra-fast components (i.e. **Figure 2B**). Bi-Gauissian curves were used to fit the distribution of tau and the areas under each of the two Gaussian curves corresponding to the ultra-fast and fast endocytosis events were determined. The ultrafast endocytosis ratio were defined by dividing the area corresponding to the ultrafast component by the sum of ultrafast and fast areas.

To determine the fraction of retrieval, event traces were fitted using a mono-exponential curve. The fraction of retrieval was determined by dividing the difference of the amplitude values of the curve at peak and at the end of the 1sec period following the stimulation by the value at the event’s peak amplitude. Calculated fractions of retrieval were categorized into four groups: “Partial” and “excessive” retrieval corresponded to fractions <0.8 or >1.2, respectively. Fractions between 0.8-1.2 were categorized as “quantal”, and “no retrieval” was determined if no detectable signal decay was observed during the 1 sec period.

The dwell time curves were determined using an exponential decay curve after a specific period of a plateau. A combination of plateau and decay curves was used for fitting the endocytosis data. Dwell time and half-time were calculated from the fits by determining the plateau periods before the decay and time to reach half of the peak values.

#### Synapse colocalization with mitochondria

The MitoTracker™ Red FM was imaged using 575 nm LED (with a filter cube containing T580lpxr, ET685/70) and recorded for 50 seconds at 50ms/frame. Synaptic activity was recorded using imaging of VGluT1-pHluorin using a near-TIRF approach described above. Particle tracking showed that only ∼2% of mitochondria exhibited fast mobility (>0.5µm/s), while the vast majority were nearly immobile (∼80% had speeds <0.1µm/s). Average images of all frames in each of the two channels were created to use for co-localization analysis. First, for mitochondria localization in the red channel, the smaller noise components below 175 nm were removed from the average image using a smaller removal filter. Subsequently, a binarized mask was generated by computing locally adaptive thresholds, which were selected based on local image statistics around each pixel. Objects exceeding a diameter of 5 μm were eliminated from consideration. The threshold intensity of 100 AU was determined based on the average intensity of the axons, excluding the mitochondria and detections with an intensity below the threshold were excluded from analysis. The resulting binarized mitochondrial localization was then utilized to compute the distance from the center of each synapse to the closest edge of the mitochondria. Areas near dense neurite intersections when it was uncertain whether the mitochondria and synapse belonged to the same axon were excluded. Subsequently, synapses were categorized based on the threshold distance of 1 μm between the bouton center and the nearest mitochondria edge into two groups: those with mitochondria (<1 μm)_and those without (>1 μm).

#### Analysis of cytosolic calcium dynamics

Cytosolic calcium measurements in response to 50AP at 40Hz or 10AP at 1 Hz were recorded in 5mM glucose, or 1.5mM Lactate and 1.5mM Pyruvate. Neurons were imaged at a frame rate of 10Hz before and 5 minutes after application of either 2μm oligomycin, or 5mM 2-Deoxy-glucose. As a standard, 50 frames were recorded before the stimulus train was triggered. The fluorescence intensity of about 20 nerve terminals (from a single neuron) was determined using the Time Series Analyzers plugin in ImageJ. Background fluorescence was subtracted from all fluorescence values and the change in fluorescence (ΔF) was obtained by subtracting the average fluorescence of the first 50 frames (F_0_). ΔF/F_0_ values were then plotted, and peak values and decay rate were determined. Fluorescence decay rate (tau) was calculated by fitting the fluorescent change after the stimulus to a single exponential decay. Peak fluorescence of 50AP trains at 40Hz was calculated by averaging the ΔF/F_0_ values of the last 5 frames of stimulus train. Single AP responses were obtained by averaging the ΔF/F_0_ values of 10 APs triggered at 1Hz. At least 10 neurons from multiple coverslips and cultures were recorded and responses were averaged.

#### Data Inclusion and Exclusion Criteria

A minimum of 10 detected release events per bouton was required for all spatial and temporal analyses of exo- and endocytosis to ensure adequate determination of AZ dimensions and adequate sampling of event spatiotemporal features across individual AZs. For all experiments, including the determination of Pr, functional synapses were defined as those with a Pr ≥0.05; synapses with a Pr below this threshold were excluded from analysis. Exclusion criteria for synapse/mitochondria co-localization analysis are described in the corresponding analysis section.

#### Statistical analysis

Statistical analyses were performed in Matlab. Statistical significance was determined using a two-sample two-tailed t-test, Tukey-Kramer ANOVA, and Kolmogorov-Smirnov(K-S) tests where appropriate. Statistical tests used to measure significance, the corresponding significance level (p-value), and the values of n are provided for each panel in **Table S1**. Data are reported as mean ± SEM, and p < 0.05 was considered statistically significant.

## SUPPLEMENTARY FIGURE LEGENDS

**Supplementary Figure 1. The effects of metabolic inhibition on spatial distribution of synaptic vesicle release and intra-terminal calcium dynamics.**

**(A,C)** Cumulative distribution of the distances between release events within individual synapses maintained in solution containing glucose or pyruvate/lactate **(A)**, or 2DG **(C)**, with or without oligoM.

**(B,D)** Average distances between release events within individual synapses in data from (**A**) or (**C**).

**(E-G)** Average traces of intra-terminal calcium response to 1 AP stimulation in neurons expressing jGCaMP8f in solutions containing glucose **(E)**, pyruvate/lactate **(F)**, or 2DG **(G)**, with or without oligoM.

**(H)** Amplitudes of calcium response to single AP stimulation under different metabolic conditions.

**(I)** Average decay constant (tau) of calcium response following stimulation.

***p < 0.001, n.s., not significant. K-S test or paired t-test.

**Supplementary Figure 2. Metabolic vulnerabilities of fast and ultrafast endocytosis.**

**(A,C)** Average event traces evoked by 1 AP stimulation and imaged in glucose or pyruvate/lactate solution **(A)**, or 2DG **(C)**, with or without mitochondrial inhibitors OligoM and UK5099.

**(B,D)** Distribution of endocytosis tau values determined from mono-exponential fits of individual event decays in different metabolic conditions.

**(A)** Representative average traces of all events evoked by 1 AP stimulation in a single synapse. Exponential decay fitting curves are indicated in red.

**(B)** Distribution of endocytosis tau values of the event traces averaged separately for each synapse.

**(C)** Ratio of ultrafast endocytosis as determined for events averaged within individual synapses under different metabolic conditions.

**(H,I)** Cumulative comparison of fraction of retrieval in glucose or pyruvate/lactate solutions **(H)**, or 2DG **(I)**, with or without UK5099 or oligoM.

** < 0.01, ***p < 0.001, n.s., not significant. One-way ANOVA.

**Supplementary Figure 3. Metabolic dependency of endocytosis dwell time and half-time.**

**(A)** Representative traces of vesicle endocytosis with varying dwell times following 1 AP stimulation.

**(B-D)** Distribution of dwell times of release events in glucose or pyruvate/lactate solutions, or 2DG with or without oligoM or UK5099.

**(E-G)** Distribution of event decay half-time, defined as the time required to reach half of the peak intensity after 1 AP stimulation under the same metabolic conditions as (B-D).

***p < 0.001, n.s., not significant. K-S test.

**Supplementary Figure 4. The impact of metabolic inhibition on temporal properties of univesicular and multi-vesicular release.**

**(A)** Example of vGlut1-pH recording with identified UVR (black) and MVR (red) release events in a single bouton, along with corresponding images. Scale bar: 1 μm.

**(B)** Distribution of the amplitudes of all detected release events from singe synapses with two peaks corresponding to UVR (q1) and MVR (q2), along with their respective Gaussian fits in light and dark blue, respectively.

**(C)** Probability of MVR under different metabolic conditions.

**(D,F)** Distributions of endocytosis tau values following UVR (light blue) and MVR (dark blue) events. Inset: ultrafast endocytosis ratio of two modes.

**(E,G)** Cumulative comparison of endocytosis tau values for UVR and MVR.

**(H-I)** Ratio of ultrafast endocytosis following UVR **(H)** and MVR **(I)** under different metabolic conditions.

** < 0.01, ***p < 0.001, n.s., not significant. One-way ANOVA.

## Notes

### Competing Interest Statement

The authors have declared no competing interest.

