## Supplementary figures and images for "Metabolic Regulation of Single Synaptic Vesicle Exo- and Endocytosis in Hippocampal Synapses"

### Supplementary Figures

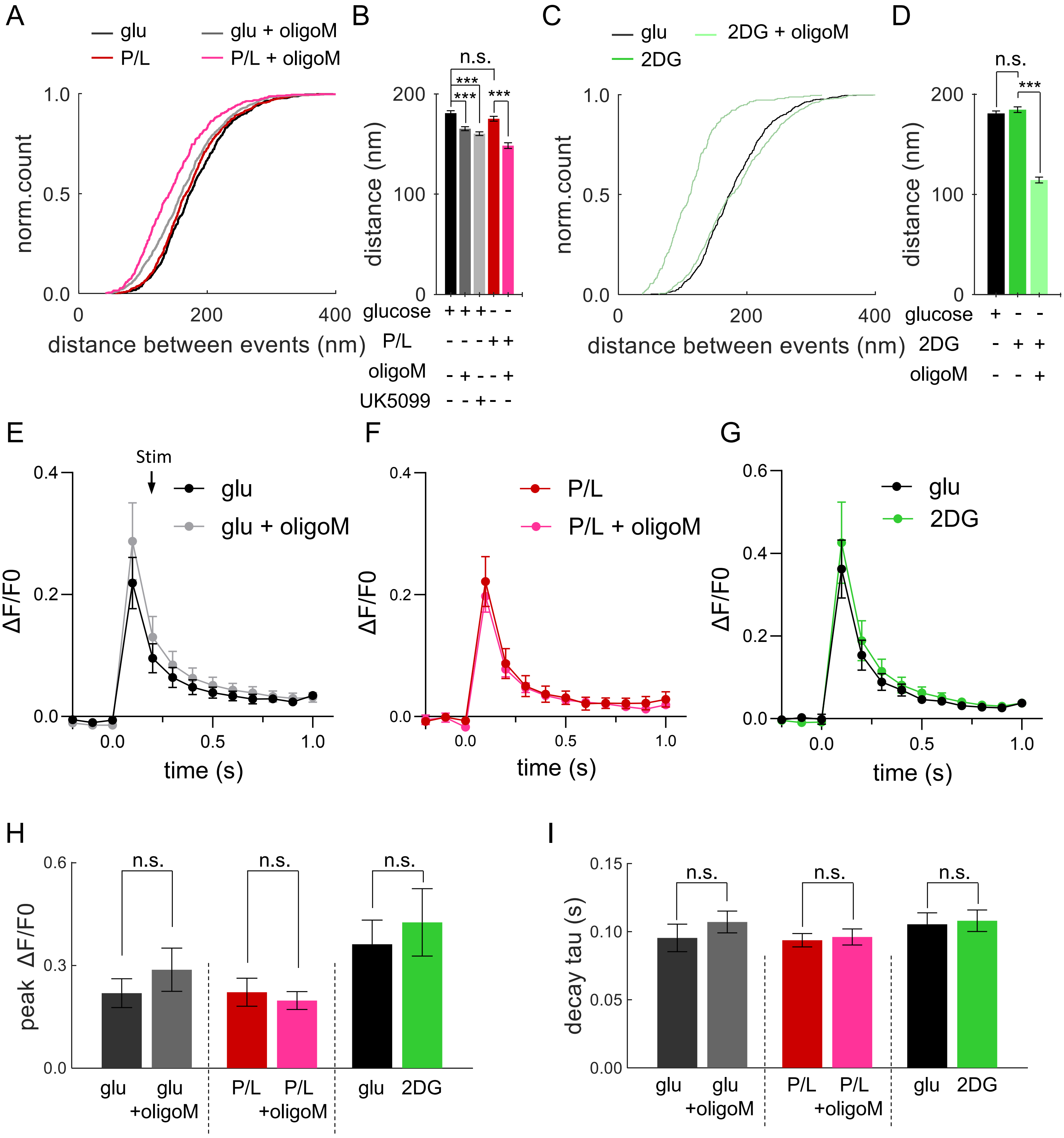


Supplementary Figure 1

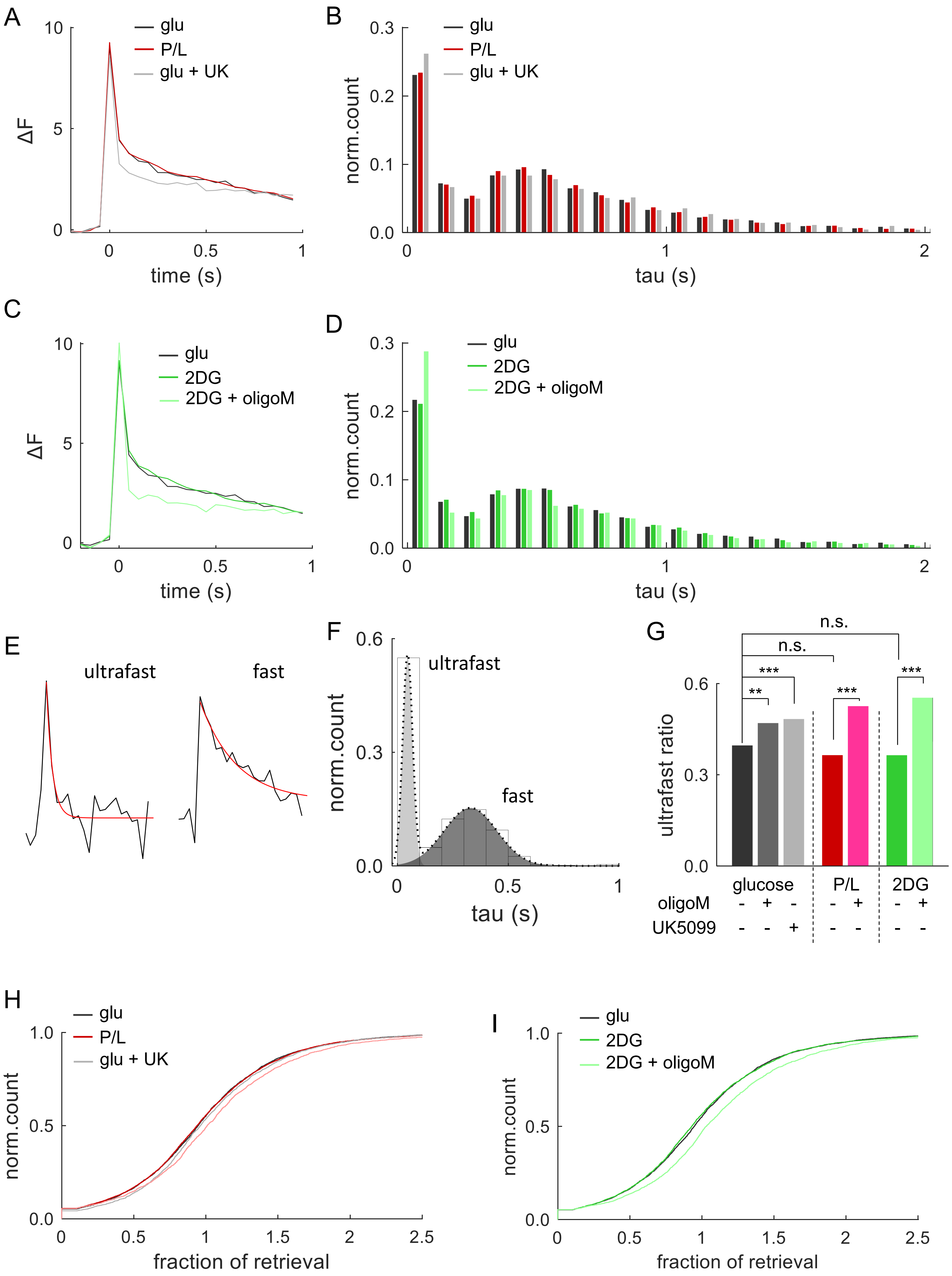


Supplementary Figure 2

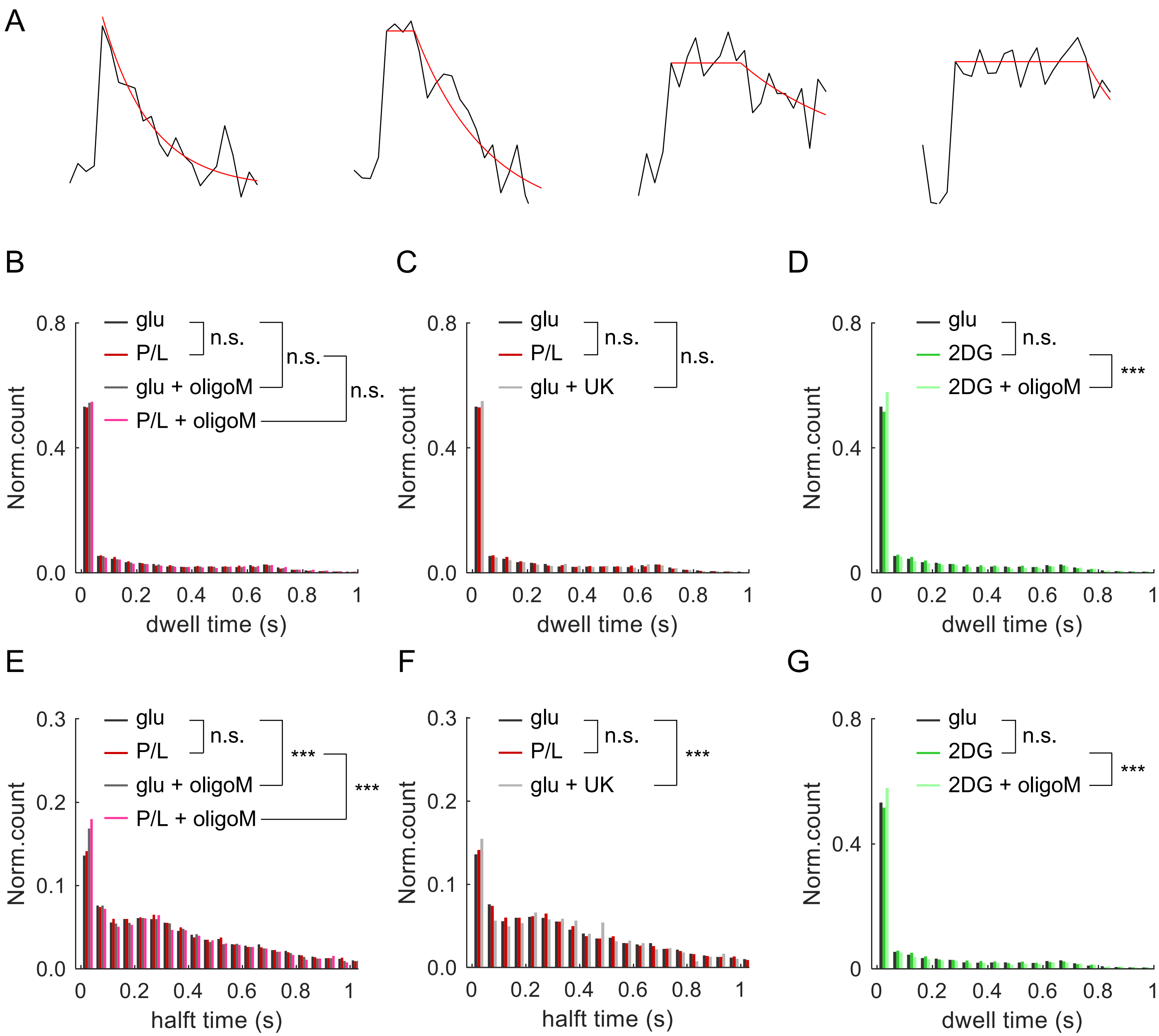


Supplementary Figure 3

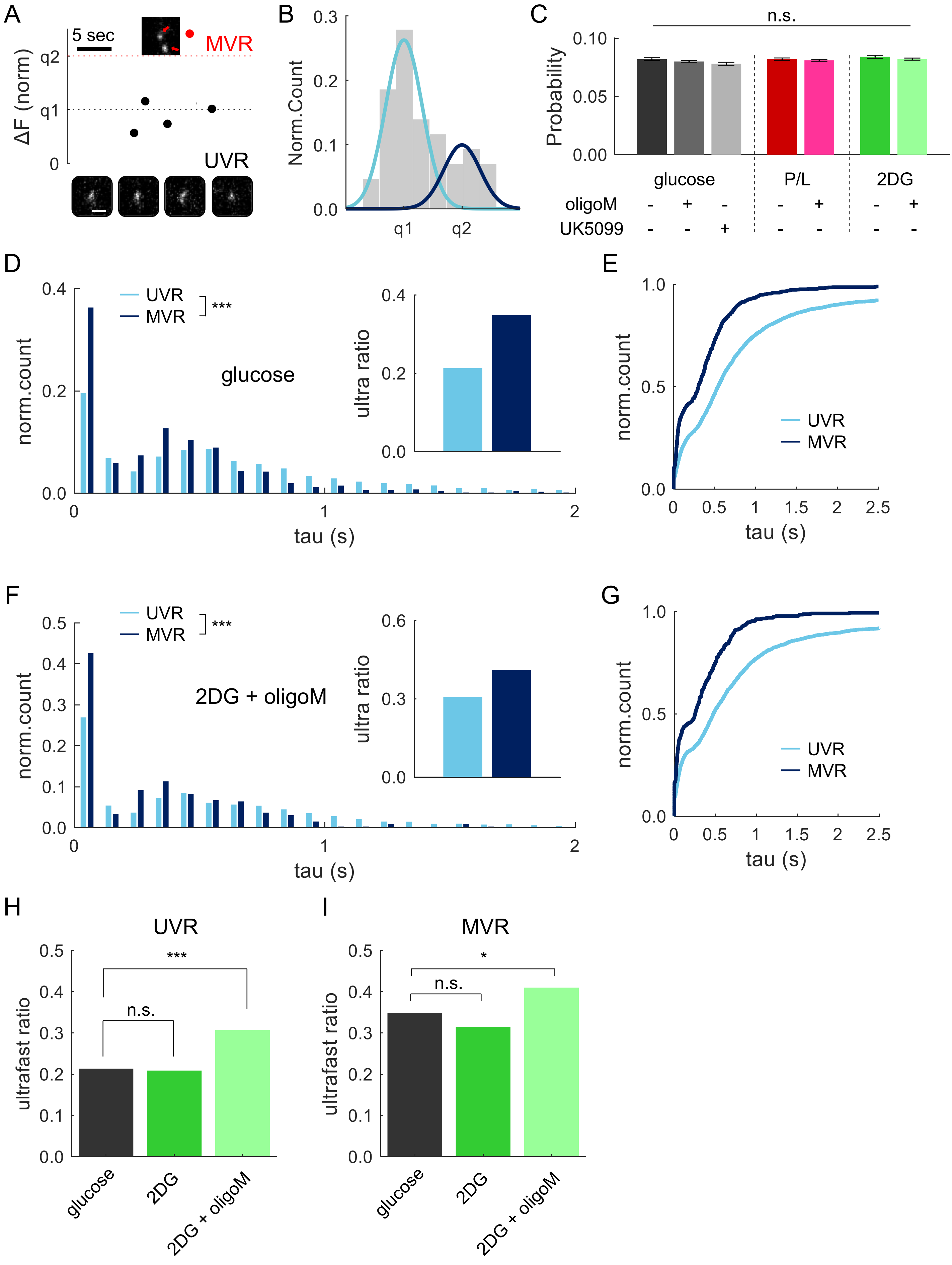


Supplementary Figure 4
